# CryoET shows cofilactin filaments inside the microtubule lumen

**DOI:** 10.1101/2023.03.31.535077

**Authors:** Camilla Ventura Santos, Stephen L. Rogers, Andrew P. Carter

## Abstract

Cytoplasmic microtubules are tubular polymers that can harbor small proteins or filaments inside their lumen. The identity of these objects and what causes their accumulation has not been conclusively established. Here, we used cryogenic electron tomography (cryoET) of *Drosophila* S2 cell protrusions and found filaments inside the microtubule lumen, which resemble those reported recently in human HAP1 cells. The frequency of these filaments increased upon inhibition of the sarco/endoplasmic reticulum Ca^2+^ ATPase (SERCA) with the small-molecule drug thapsigargin. Subtomogram averaging showed that the luminal filaments adopt a helical structure reminiscent of cofilin-bound actin (cofilactin). Consistent with this, cofilin was activated in cells under the same conditions that increased luminal filament occurrence. Furthermore, RNAi knock-down of cofilin reduced the frequency of luminal filaments with cofilactin morphology. These results suggest that cofilin activation stimulates its accumulation on actin filaments inside the microtubule lumen.

## Introduction

Microtubules are cytoskeletal filaments that provide mechanical stability and serve as tracks for cargo transport. They consist of α and β-tubulin dimers that assemble into multiple protofilaments, and come together to form the polar, tube-shape filament.

The lumen of the microtubule provides a sheltered, approx. 15 nm wide space that can accommodate small proteins (Nogales *et al*., 1999). Electron microscopy studies indeed revealed the presence of distinct, globular densities inside the microtubule lumen of neurons, whose identity and functional role remain unclear (Burton, 1984; Garvalov *et al*., 2006; Bouchet-Marquis *et al*., 2007; Atherton *et al*., 2021; Chakraborty *et al*., 2022; Foster *et al*., 2022).

Recently, a cryogenic electron tomography (cryoET) study on small-molecule induced protrusions of human HAP1 cells reported the presence of actin-based filaments inside the microtubule lumen (Paul *et al*., 2020). These luminal filaments displayed non-canonical actin morphologies, suggesting that they may be bound by further, unidentified factors. In addition to their unknown composition, microtubule luminal filaments have not been described in cells without induced protrusions, raising the question under which conditions they appear.

CryoET has been used to characterize objects inside the microtubule lumen. Thin (< 200 nm) samples generate higher signal-to-noise ratios, allowing for subtomogram averages at higher resolution. The model organism *Drosophila melanogaster* provides a set of well-characterized cell types many of which are smaller and thinner than their mammalian counterparts (Cherbas and Gong, 2014). Schneider 2 (S2) cells are derived from the *Drosophila* immune system, are highly susceptible to expression or knock-down of proteins and have been used to study the microtubule and actin cytoskeleton (Schneider, 1972; Rogers *et al*., 2003; Rogers and Rogers, 2008). They form microtubule-rich protrusions when treated with low concentrations of the actin polymerization inhibitor Cytochalsain D (CytD) (Lu *et al*., 2013), thereby providing an accessible target for microtubule-based studies with cryoET.

Using the particularly thin, CytD-induced protrusions of *Drosophila* S2 cells, we set out to answer (1) whether luminal filaments appear inside S2 cell protrusions and are therefore conserved across species, (2) what the filaments are composed of and (3) what causes their occurrence. We found rare occasions of luminal filaments in S2 cell protrusions and increased their frequency by adding thapsigargin (TG), a small-molecule inhibitor of the SERCA. Subtomogram averaging suggests that the luminal filaments are formed of cofilin-bound actin (cofilactin). In agreement with this, dsRNA mediated depletion of cofilin lead to a change in morphology of luminal filaments. Taken together, our study suggests luminal cofilactin filaments can occur inside microtubules of induced protrusions in response to specific stress.

## Results and Discussion

### CryoET of induced protrusions of *Drosophila* S2 cells

We incubated *Drosophila* S2 cells with low concentrations of CytD to induce the formation of thin protrusions (Lu *et al*., 2013). Immunofluorescence staining for α-tubulin showed that the protrusions were filled with microtubules (Fig. EV1A). We replicated this workflow on electron microscopy (EM) grids, vitrified the samples and acquired tilt series on the thinnest parts (Fig. 1A, B). We obtained tomograms in which we could unambiguously identify individual organelles and protein complexes such as the endoplasmic reticulum (ER) and ribosomes (Fig. 1C). The imaged protrusions contained parallel arrays of up to 20 microtubules and were on average 150 nm thick (Fig. EV1B – D).

**Figure 1.**
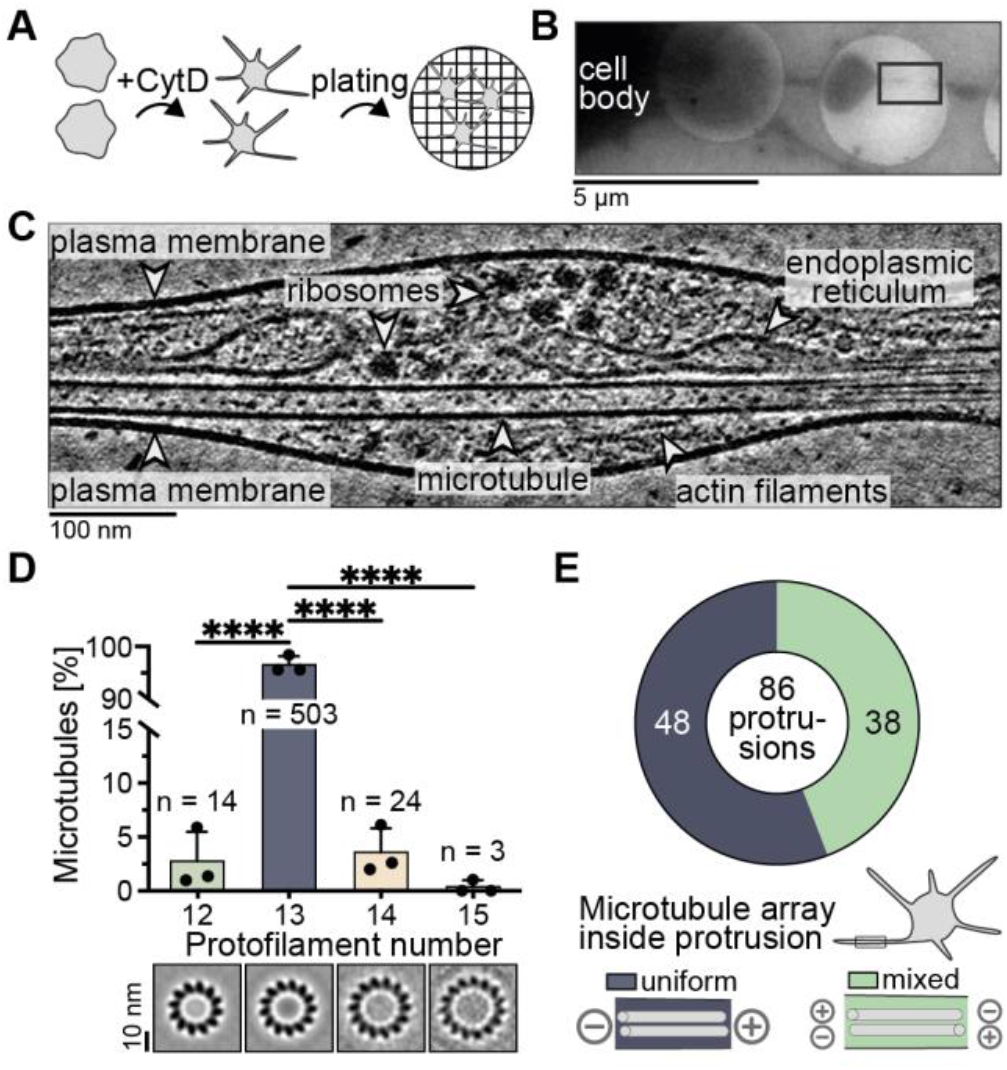
CryoET of microtubule-rich S2 cell protrusions. A) Cartoon showing cellular sample preparation for cryoET. S2 cells were mixed with CytD and then plated on Concanavalin A-coated cryoEM grids. Overview image of a vitrified sample and the area in which the tomogram in C) was acquired (black box). C) Tomogram slice of an S2 cell protrusion with different cytoskeletal filaments, organelles and protein complexes. D) Percentage of microtubules with 12 (2.8 ± 2.7%, mean ± SD), 13 (93.3 ± 3.2%), 14 (3.6 ± 2.2%) or 15 (0.3 ± 0.6%) protofilaments determined by subtomogram classification of 544 microtubules. Statistical significance was determined with an ordinary ANOVA test with multiple comparisons, only significant results are labelled with stars (p < 0.0001, ****). Bottom panels show projections of microtubule averages with different protofilament numbers. E) Pie chart of the number of protrusions with uniform (dark blue) or mixed (green) microtubule orientations determined from subtomogram classification. Only protrusions with at least 2 microtubules of resolved orientation were included (86/ 111). Cartoons at the bottom show examples of uniform and mixed microtubule arrays.

Subtomogram classification of 544 microtubules from the protrusions revealed that the majority (93.3%) had 13 protofilaments whereas minor fractions had 12 (2.8%), 14 (3.6%) or 15 (0.3%) protofilaments (Fig. 1D, EV1E). In 25 out of 86 cases, we found microtubules with different protofilament numbers within the same protrusion (Fig. EV1F). We previously showed that microtubules from *Drosophila* neurons can have 12 or 13 protofilaments and coexist within the same neurite (Foster *et al*., 2022). Our results from S2 cells provide another example of a cell type in which different protofilament architectures can coincide within the same cell.

We next used a similar classification approach to analyze the orientations of the microtubules inside each protrusion (Sosa and Chrétien, 1998; Bouchet-Marquis *et al*., 2007; Foster *et al*., 2022) (Fig. EV1E). We analyzed protrusions in which we could resolve the polarity of all microtubules. These contained between 2 and 16 microtubules per protrusion. We found 48 out of 86 contained a uniform array of microtubules (Fig. 1E). The remainder contained between 1 and 6 microtubules with opposite orientation compared to the majority. In this analysis, we observed a variation across biological replicates, with two datasets showing ∼46% uniform orientation and one dataset where 17 out of 19 protrusions had microtubules pointing in the same direction (Fig. EV1G).

Studies using the fluorescently tagged, tip-tracking protein EB1, found that 80 – 95% of microtubules are uniformly oriented inside S2 protrusions (Kural *et al*., 2005; del Castillo *et al*., 2015; Del Castillo *et al*., 2020). Together with our data this suggests that many, but not all, *Drosophila* S2 cell protrusions contain uniformly oriented microtubules.

### Filaments inside the microtubule lumen increase upon inhibition of the SERCA

We inspected the microtubules in our tomograms and found rare instances of filaments inside their lumen (Fig. 2A). These filaments had similar appearance to filamentous actin (f-actin) found in the microtubule lumen of human HAP1 cells (Paul *et al*., 2020).

**Figure 2.**
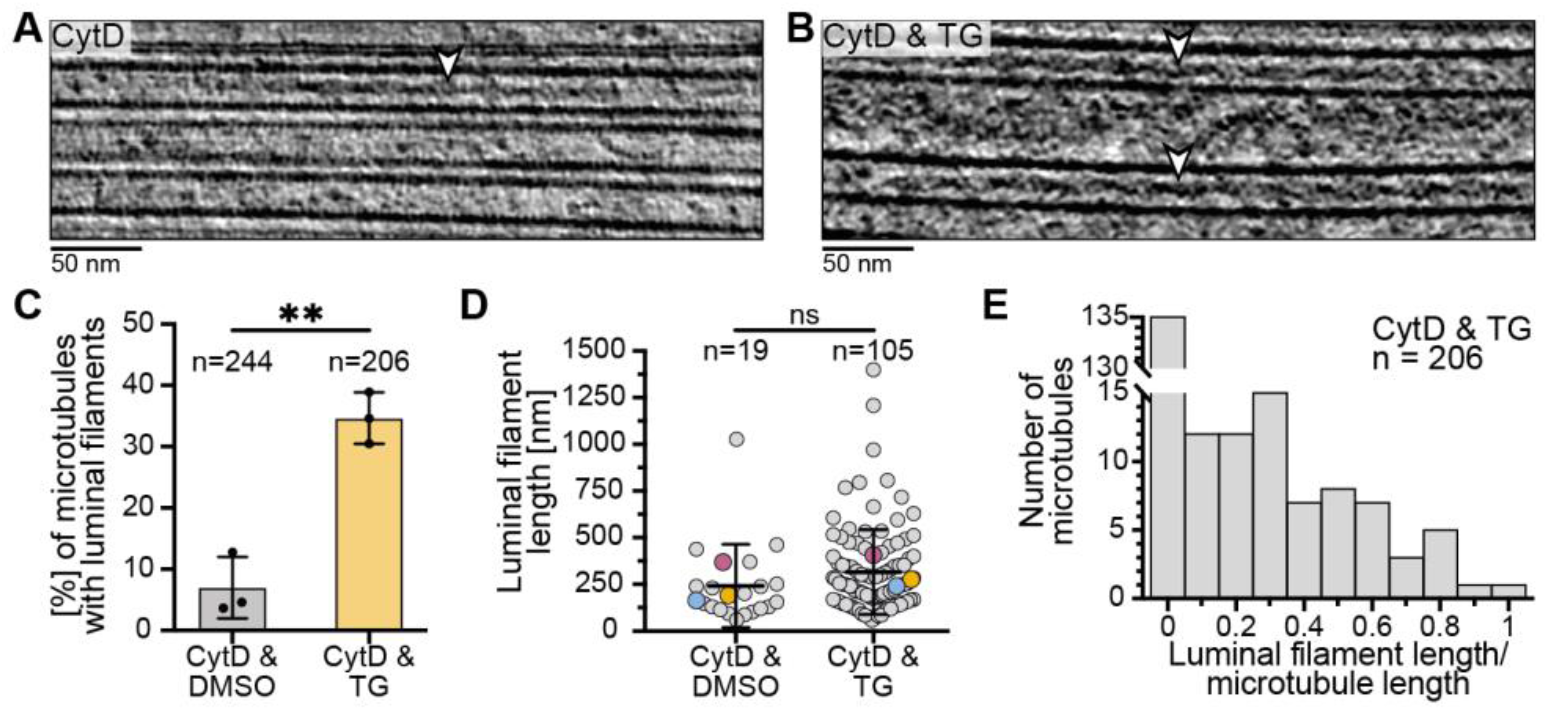
SERCA inhibition with TG increases occurrence of luminal filaments. A, B) Tomogram slices showing filaments inside the microtubule lumen (white arrowheads) of cells treated with CytD (A) or with CytD & the SERCA inhibitor TG (B). C) Percentage of microtubules containing at least one luminal filament in cells treated with CytD & DMSO (7.0 ± 5.0%, mean ± SD) or CytD & TG (34.7 ± 4.2%) from three biological replicates. Statistical significance was determined with an unpaired t-test (p = 0.0019, **). D) Length of luminal filaments from cells treated with CytD & DMSO (241.8 ± 110.9 nm, mean ± SD) or CytD & TG (308.1 ± 87.1 nm). The means of three biological replicates are shown in colored circles and were used for a non-parametric Mann-Whitney test (p = 0.4, ns). E) Histogram of the fraction of microtubule length that is encompassed by luminal filaments for each microtubule in cells treated with CytD & TG. The total length of luminal filaments in one microtubule was divided by the length of that microtubule. 71 out of 206 analyzed microtubules from cells treated with CytD & TG contained at least one luminal filament. Of these, the majority (54.9%) had a length coverage of 0.1 - 0.3.

Luminal filaments were too rare for an analysis of their distribution and structure and so we set out to increase their frequency. We hypothesized that the filaments may form in response to specific cell stresses and tested if addition of a drug increased their occurrence. We tested small-molecule inhibitors of the proteasome (MG132) or the SERCA (TG) to disturb protein and Ca^2+^ homeostasis, respectively (Thastrup *et al*., 1990; Lee and Goldberg, 1998; Wu *et al*., 2002; Lu *et al*., 2014). While there was no increase in luminal filaments upon proteasome inhibition, SERCA inhibition increased their occurrence in a preliminary experiment (Fig. EV2A). We therefore prepared samples for cryoET from three biological replicates by treating cells with CytD and either a vehicle (DMSO) or TG (Fig. 2B, EV2B). We acquired tilt series of these samples and found 105 luminal filaments in TG treated cells. In contrast, only 19 luminal filaments were present in cells treated with a vehicle. We analyzed how many microtubules contained luminal filaments. 7.0 ± 5.0% of microtubules had luminal filaments in DMSO treated cells (Fig. 2C). In contrast, treatment with TG significantly increased this value 5-fold to 34.7 ± 4.2% (Fig. 2C). Taken together, we observed filaments inside the microtubule lumen whose frequency increased upon inhibition of the SERCA with TG.

The filaments had varying lengths with the majority (75%) ranging from 58 – 400 nm. There was no significant difference between control and TG treated cells (Fig. 2D), suggesting that SERCA inhibition increases the frequency but not the length of luminal filaments. Our analysis of the filaments in cells treated with TG showed that most of them only spanned a fraction of the microtubule length (Fig. 2E). In rare cases, microtubules contained 2 and up to 4 filaments inside their lumen (Fig. EV2C, D), consistent with their short length compared to the microtubules.

The luminal filaments occupied a large volume of the microtubule lumen and we asked if this affected the ultrastructure of the surrounding microtubule, for example by favoring microtubules with larger diameters. We analyzed the protofilament number distribution and found no difference between microtubules with and without luminal filaments (Fig. EV2E). We even found a single case where a luminal filament was present in a 12 protofilament microtubule. These results suggest that luminal filaments do not affect the structure of the surrounding microtubule.

### Structural analysis suggests that luminal filaments are formed of cofilactin

We inspected the luminal filaments using Fourier transforms to identify repeating units and their frequencies. The luminal filaments periodically widened and compacted every 27 ± 0 nm (Fig. 3A, EV3A). This so-called cross-over distance was significantly shorter than the ∼36 ± 2 nm repeat determined for cytoplasmic f-actin (Fig. 3B, EV3B, C). The luminal filaments also had wider diameters than cytoplasmic actin (Fig. 3A, B), consistent with the hypothesis that they are formed of actin and additionally bound proteins.

**Figure 3.**
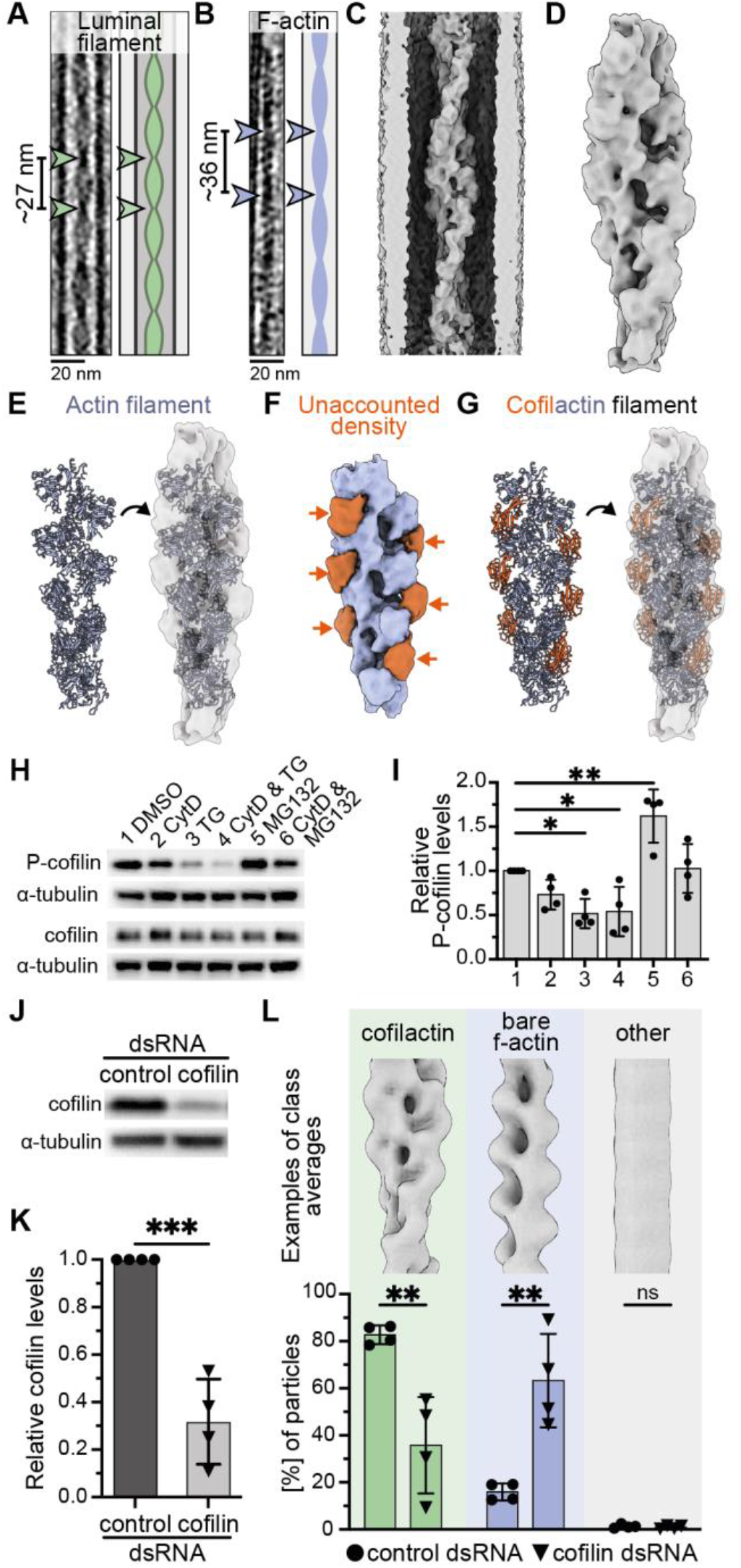
Subtomogram averaging of luminal filaments suggests they are formed of cofilactin. A, B) Tomogram slices and cartoons of a luminal filament (A) and cytoplasmic f-actin (B). The cross-over distances derived from Fourier transforms (Fig. EV3A – C) are indicated (arrowheads). C, D) Unmasked (C) and masked (D) subtomogram average structure of the luminal filament at 16.5 Å. E) Fit of an actin filament into the subtomogram average density shown in D). F) Density representation showing ‘unaccounted density’ from E) in orange. G) Fit of a cofilactin model (actin: blue, cofilin: orange) into the subtomogram average structure of the luminal filament. H) Western blot showing that TG decreases phosphorylated cofilin (P-cofilin). Top: P-cofilin, bottom: total cofilin after treatment of cells with DMSO, CytD, TG or MG132. α-tubulin was used as loading control. I) Quantification of western blots from the experiment shown in H (number labels are as in H): DMSO (1 ± 0, mean ± SD), CytD (0.73 ± 0.17), TG (0.52 ± 0.16), CytD & TG (0.54 ± 0.28), MG (1.62 ± 0.30), CytD & MG (1.03 ± 0.28). An ordinary ANOVA test with comparisons to the DMSO control was performed (DMSO vs. TG, p = 0.0276, *; DMSO vs. CytD & TG, p = 0.0370, *; DMSO vs. MG, p = 0.0043, **, other comparisons, p > 0.05, ns). J, K) Western blot (J) and quantification (K) showing that cofilin protein levels were reduced to 31.8 ± 18.0% on day 7 or 8 of dsRNA treatment in S2 cells. L) Top: Examples of class averages of different luminal filament types. Bottom: Percentages of luminal filament particles classified into cofilactin (green, 82.7 ± 3.9% control dsRNA, 35.8 ± 20.5% cofilin dsRNA, mean ± SD), bare f-actin (blue, 15.9 ± 3.6% control dsRNA, 63.3 ± 19.9% cofilin dsRNA) or other (grey, 1.3 ± 0.8% control dsRNA, 0.9 ± 0.7% cofilin dsRNA) classes. Unpaired t-tests were used to compare classifications from control dsRNA (circles) and cofilin dsRNA (triangles) samples. P-values are: cofilactin (0.0041, **), bare f-actin (0.0034, **) and other (0.4784, ns).

We applied subtomogram averaging to gain structural information of the luminal filaments. To acquire sufficient particles, we combined luminal filaments from multiple datasets (see Materials and methods). We manually picked particles, performed initial alignments in Dynamo (Castaño-Díez *et al*., 2012), and then classified and refined particles in Relion 3 (Scheres, 2012; Zivanov *et al*., 2018) (Fig. EV3D). Averaging of individual filaments showed that they were polar (Fig. EV3D). We therefore determined the polarity of each filament and flipped particles accordingly to obtain a uniformly oriented particle set. Alignment of the luminal filaments lead to smearing of the surrounding microtubule density and vice-versa indicating that the luminal filaments were not regularly positioned with respect to the microtubule (Fig. EV3E). This misalignment made the filaments a challenging target for subtomogram averaging.

Our structure at 16.5 Å revealed a helical filament inside the microtubule lumen (Fig. 3C, D, EV3F). We first fit an actin filament into our structure and found repeating densities on the edges of the filament that were unaccounted for (Fig. 3E, F). We noticed that the helical twist, which describes the angle between two neighboring subunits around the filament axis, was reduced in the luminal filament compared to bare f-actin. To identify actin-binding candidates that could induce a change in helical symmetry and bind at the position of the unaccounted density, we searched the electron microscopy database (EMDB) for helical filaments containing actin. Bare actin and actin bound to different families of actin binding proteins had consistent helical parameters with a helical twist of approx. -167° (Appendix Table S1). Only entries of cofilin-bound actin (cofilactin) had a reduced helical twist of -162° (McGough *et al*., 1997; Tanaka *et al*., 2018) and this value matched the parameter determined for our luminal filament structure (−161.8°). In agreement with this, a cofilactin model fit well into our structure and the cofilin moiety occupied the extra density that had been unaccounted for when fitting actin alone (Fig. 3G).

### Cofilin is activated upon SERCA but not proteasome inhibition

Cofilin is a key regulator of the actin cytoskeleton and highly conserved from yeast to humans with a single homologue in *Drosophila* called *twinstar* (Edwards *et al*., 1994; Gunsalus *et al*., 1995). The affinity of cofilin for actin is strongly increased upon dephosphorylation at a conserved site (Ser 3) and cells use this post-translational modification to control cofilin’s actin binding and severing activity (Agnew *et al*., 1995; Zebda *et al*., 2000; Austin Elam *et al*., 2017).

Our structural analysis suggested that luminal filaments were formed of cofilactin (Fig. 3G) and the frequency of luminal filaments was increased upon treatment of cells with TG (Fig. 2C). We asked if TG activates (i.e. dephosphorylates) cofilin as this could explain the observed increase in luminal cofilactin filaments. To test this, we treated S2 cells with CytD, TG or MG132 and used antibodies against phosphorylated cofilin (P-cofilin) in western blots. Indeed, TG treatment decreased P-cofilin, indicating cofilin activation (Fig. 3H, I). In contrast, MG132 deactivated cofilin by increasing its phosphorylation (Fig. 3H, I). These results are consistent with our findings from cryoET where TG but not MG132 treatment increased the frequency of luminal filaments (Fig. 2C, EV2A), and provide further support for a role of cofilin in luminal filament formation.

### Luminal filaments have altered morphology in cofilin knock-down cells

To gain further evidence for the presence of cofilin in luminal filaments, we sought to deplete S2 cells of cofilin. We used a double-stranded RNA (dsRNA) to reduced cofilin levels to 31.8 ± 18.0% compared to a control dsRNA (Fig. 3J, K). We then induced the formation of protrusions and luminal filaments with CytD & TG and prepared samples for cryoET. To determine changes in luminal filament morphology, we performed subtomogram classification and quantified the results (Fig. EV3G, H). In cells treated with a control dsRNA, 82.7 ± 3.9% of luminal filament particles fell into classes with cofilactin morphology (Fig. 3L). In contrast, in cofilin knock-down cells this value was reduced to 35.8 ± 20.49% and instead 63.3 ± 19.9% of particles were present in classes resembling bare f-actin (Fig. 3L). We obtained similar results from visual inspection of filament morphologies in the tomograms (Fig. EV3I). A small fraction of luminal filaments was neither cofilactin nor bare f-actin and adopted alternative filament morphologies (Fig. EV3J). The increase in bare f-actin within the microtubule in the absence of cofilin provides direct evidence that the luminal filaments we observed contain cofilin.

In summary, we combined cryoET, subtomogram averaging and RNA interference to show that the filaments we observed inside the microtubule lumen of *Drosophila* S2 cells are formed of cofilactin. Furthermore, we provide evidence that these filaments specifically accumulate in response to TG. Although this SERCA inhibitor induces ER stress, we suspect that luminal filament accumulation is not a general stress response as we did not observe it in the presence the proteasome inhibitor MG132. Instead, we suggest luminal filament accumulation is due to TG directly activating cofilin in S2 cells.

Cofilin is an actin binding protein that severs f-actin at low concentrations and decorates it to form cofilactin at high concentrations (Nishida *et al*., 1984; Maciver *et al*., 1991; Andrianantoandro and Pollard, 2006). It is activated both by cellular stresses, such as energy depletion (Nishida *et al*., 1987; Meberg *et al*., 1998; Minamide *et al*., 2000), and by signaling pathways, particularly those found at the leading edge of the cell (Arber *et al*., 1998; Meberg *et al*., 1998; Yang *et al*., 1998; Chan *et al*., 2000; Zebda *et al*., 2000; Endo *et al*., 2003). Indeed, filaments resembling cofilactin have been observed in the cytoplasm of neuronal growth cones in two recent cryoET studies (Atherton *et al*., 2022; Hylton *et al*., 2022). It remains to be seen whether cofilactin can also accumulate in the microtubule lumen during cell migration.

Although cofilin knock-down in S2 cells lead to a major change in the filament morphology, we observed only a minor (and non-significant) decrease in the total frequency of luminal filaments (Fig. EV3K). We speculate this may be due to the residual cofilin remaining in the cells (Fig. 3J, K) being able to nucleate filaments inside the microtubule. This would be consistent with our observations of luminal filaments that transitioned from a cofilactin to a non-cofilactin morphology (Fig. EV3L).

It is unclear how actin and cofilin get inside the microtubule. Entry of preassembled cofilactin filaments via microtubule ends appears unlikely given the narrow diameter of the microtubule lumen in comparison to cofilactin filaments. Individual components could enter through breaks in the microtubule lattice (Chakraborty *et al*., 2020; Atherton *et al*., 2022; Foster *et al*., 2022). Alternatively, microtubules may polymerize around pre-existing cytoplasmic filaments, resulting in their encapsulation. In this case, actin filaments could “guide” the polymerization of microtubules thereby influencing organization and dynamics of the microtubule network.

The physiological role of cofilactin within the microtubule lumen, if any, remains unknown. One possibility is that they stabilize the surrounding microtubule. In our best tomograms we observed connections between luminal filaments and the microtubule wall (Fig. EV3M), which could structurally reinforce the microtubule lattice. A similar stabilizing role has been proposed for the globular luminal particles found in neurons and these are also frequently linked to the surrounding microtubule via short tethers (Garvalov *et al*., 2006; Cuveillier *et al*., 2020; Chakraborty *et al*., 2022; Foster *et al*., 2022). Another possibility is that uptake of cofilactin filaments into the microtubule lumen acts as a sink for excessive active cofilin. A related mechanism, forming cytoplasmic bundles of cofilactin filaments, was previously proposed as a protective mechanism for stressed cells to reduce energy consumption (Bernstein *et al*., 2006). In either case, the observation that microtubule luminal filaments were also found in human cells (Paul *et al*., 2020) suggests that their function may be conserved across species.

## Materials and Methods

### Cell culture

Schneider S2 cells were grown in Schneider’s medium (21720001, Gibco or 04-351Q, Lonza) supplemented with 10% heat-inactivated FBS, 100 units/ mL Pen/Strep and 0.25 μg/ mL Amphotericin B (15290018, Gibco) at 25°C. Cells were split every 2 – 4 days and maintained until passage 25.

### Generation of a *Drosophila* α-tubulin-acetylase (dTAT) knock-out line using CRISPR

The dTat knock-out cell line was made using protocols created by Bassett *et al*., 2014. A pair of 20-mer oligonucleotides targeting the 5’ end of the *dtat* coding sequence was ligated into pAc-sgRNA-Cas9 (Addgene) and validated by sequencing. The construct was transfected into low pass S2 cells using TransIT Insect transfection reagent (Mirus) and stable cells carrying indels for this locus were selected for with 5 μg/ ml puromycin for 3 weeks. Loss of dTAT was validated by immunoblotting and immunofluorescence for acetylated α-tubulin (Appendix Figure S2A, B). The dTAT knock-out cell line will be distributed through the Drosophila Genomics Resource Center (Bloomington, IN).

### Double-stranded RNA interference

dsRNAs were synthesized by *in vitro* transcription from PCR products using forward and reverse primers containing the T7 promoter sequence (TAATACGACTCACTATAGG) with homemade Pfu. Amplicons were selected by querying gene specific sequences using http://www.genomernai.org/ and amplified from genomic DNA isolated from S2 cells using Trizol purification. Cofilin/ *twinstarI* dsRNA was synthesized using primers (5’-TAATACGACTCACTATAGGCATTTCTGGATATCTTCTAGAAACT-3’ and 5’-TAATACGACTCACTATAGGTGGTGTAACTGTGTCTGATGTCTG-3’). PCR product sizes were validated by agarose gel electrophoresis and concentrated by precipitation with sodium acetate and cold ethanol. dsRNA was synthesized overnight at 37°C using T7 RNA polymerase (FisherBiosci) in transcription buffer (40 mM Tris-HCl pH 8.0; 10 mM DTT; 2 mM Spermidine-HCl (Sigma); 20 mM MgCl2; 7.5 mM rNTPs (Promega); inorganic pyrophosphatase (FisherBiosci)). *In vitro* transcription reactions were treated with DNAse (NEB) for 1 hour at 37°C, quantitated for yield by comparison against a DNA ladder, precipitated with sodium acetate and cold ethanol, and resuspended in DNAase/ RNAse-free water to a concentration of 1 mg/ mL. For all experiments, we included a negative control with dsRNA corresponding to the sequence of bacterial chloramphenicol transferase using primers (5’-TAATACGACTCACTATAGGGATCCCAATGGCATCGTAAAGAACATTTGAGGC-3’ and 5’-TAATACGACTCACTATAGGGGGGCGAAGAAGTTGTCCATATTGGCCA-3’).

For control and cofilin knock-down experiments, 1 mL of dense S2 cell culture was added to 9 mL of serum-free S2 media (Schneider’s medium, 100 units/ mL Pen/Strep) and pelleted for 10 min at 600 x g. The supernatant was removed and cells were plated at a density of 5 * 10^5^ cells/ mL in 1 mL serum-free media in a well of a 12 well plate and allowed to adhere to the surface for 20 – 30 min. The media was then replaced with 500 μL serum-free media containing 7 (replicate 1) or 20 μg (all other replicates) dsRNA for control or cofilin RNA interference and incubated for 1 h at 25°C in a humidified chamber. Then, we added 500 μL of S2 media with 20% FBS. The same procedure was repeated 3 days later. Cells were frozen on day 7 or 8 of the experiment. Knock-down efficiencies of all replicates used for cryoET data acquisition were examined by western blotting for total cofilin. Western blots and quantifications were performed as described in the section ‘Western blotting and quantification’ with the modification that only approx. 1 – 10 * 10^5^ cells were used. Cells were preincubated with 2 μM TG for 3 h prior to lysis and western blotting, because part of each batch was also used for the cryoET sample preparation.

### Western blotting and quantification

For western blots of P-cofilin, we seeded cells at 2 * 10^6^ cells/ mL in S2 media in 2 mL media/ well in a 6-well plate with either DMSO (D2650, Merck), 2.5 μM CytD (C2618, Sigma), 2 μM TG (ab147487, Abcam), 2.5 μM CytD & 2 μM TG and, 25 μM MG132 (M7449, Merck) or 2.5 μM CytD & 25 μM MG132. All samples were adjusted to 0.275% DMSO. Cells were incubated with the drugs for 5 h at 25°C, then washed with 1 mL ice-cold PBS and lysed in 70 μL RIPA buffer/ well (89900, Thermo Scientific) supplemented with cOmplete protease inhibitor cocktail (11697498001, Sigma) and Halt™ phosphatase inhibitor cocktail (1862495, Thermo Scientific). All following steps were performed at 4°C unless stated otherwise. Cells were scraped off with the tip of the pipet and transferred to a 1.5 mL tube. Samples were incubated for 15 min and then centrifuged for 12 min at 21,000 x g. Supernatants were transferred to a fresh tube and boiled with NuPAGE LDS Sample buffer (4x, NP0007, Invitrogen) for 5 min. Samples were run on a NuPAGE gel at 120 – 150 V and blotted onto a PVDF membrane using a Trans-blot Turbo Transfer Systen (Biorad) in the mixed molecular weight program.

Membranes were cut above the 25 kDa band to blot for the loading control (α-tubulin) and cofilin or P-cofilin. Membranes were blocked with 5% milk in TBST (20 mM Tris-HCl, 150 mM NaCl, 0.1% w/ v Tween20, α-tubulin and cofilin) or in 5% BSA (Sigma) in TBST (P-cofilin) for 30 – 60 min at room temperature. Membranes were incubated with primary antibodies at 4°C over-night or for 1 h at room temperature at the following dilutions: α-tubulin (sc-32293, Santa Cruz, 1:5000 in 5% milk, mouse), anti-cofilin (gift from Anna Marie Sokac, 1:5000 in 5% milk, rabbit), anti-P-cofilin (gift from Buzz Baum, 1:2000 in 5% BSA, rabbit). Membranes were washed four times for 5 min each in TBST and incubated with Polyclonal goat anti-mouse (P044701, Agilent, 1:2500) or anti-rabbit (P044801, Agilent, 1:2000) immunoglobulins/ HRP for 1 h at room temperature. Membranes were again washed four times for 5 min each with TBST and developed using ECL Select Western Blotting Detection Reagent (RPN2235, Amersham). Membranes were imaged on a Bio-Rad Gel Doc imager.

We quantified the western blots by drawing a square around each band and measuring the integrated density in ImageJ (Schindelin *et al*., 2012). Bands from P-cofilin, cofilin and α-tubulin were first normalized to the band intensity of the respective DMSO treatment. P-cofilin and cofilin were then normalized to their corresponding loading controls (α-tubulin). To determine the relative phosphorylation, normalized P-cofilin band intensities were divided by normalized cofilin intensities for each sample.

### Immunofluorescence and light microscopy imaging

Autoclaved 13 mm coverslips were placed in wells of a 24-well dish and coated with 0.25 μg/ mL Concanavalin A (L7647, Sigma) for at least 1 h and up to over-night at 37°C. Coverslips were washed twice with PBS before plating 1.5 * 10^5^ cells/ well with either DMSO or 2.5 μM CytD. DMSO concentrations were adjusted to 0.125% in both samples. Cells were grown for 5 h before fixing for 10 min in fixation buffer (3% paraformaldehyde, 0.1% Glutaraldehyde, 4% Sucrose, 0.5% Triton-X-100, 0.1 M PIPES pH 7, 2 mM EGTA, 2 mM MgCl_2_). This and all the following steps were performed at room temperature. Samples were washed 3 times with PBS, permeabilized with 0.1% Triton-X-100 in PBS for 10 min and washed again with PBS. Samples were blocked with 2% BSA (A3311, Sigma) in PBS for 30 – 60 min and then incubated with primary antibody against α-tubulin (sc-32293, Santa Cruz) for 1 h. Samples were washed with PBS and incubated with secondary Alexa Fluor 647 donkey anti-mouse (life technologies) for 1 h. Samples were mounted with mounting media (P36962, Invitrogen) and left to cure over-night before imaging on a Zeiss 880 confocal microscope equipped with a 63x/1.4 NA Oil lens. Z-stacks including the entire volume of the cell (24 slices, 0.63 μm/ pixel) were acquired and summed to generate a projection image showing all microtubules within the cell.

### Sample preparation for cryoET

Quantifoil R3.5/1 Au200 grids (Quantifoil) were glow discharged for 30 s in an Auto306 Edwards Turbo Coater at power 6 (20 mA) and then transferred to a sterile hood. From here, samples were prepared in an upside-down p10 dish, in which the lid was covered to ∼70% with a sterile piece of parafilm. 30 μL droplets of 0.25 μg/ mL Concanavalin A were placed on the parafilm in the p10 dish and one glow-discharged grid was added to each droplet with the carbon side facing up. The p10 dish was closed and placed in a humidified chamber at 37°C for 1 h – 16 h. Before plating cells, grids were washed twice with PBS. For all experiments, we plated 8,000 - 10,000 cells in 30 μL droplets per grid and incubated them in a humidified chamber at 25°C until vitrification.

Samples used for quantification and subtomogram averaging of microtubules in protrusions were plated with either 2.5 μM (datasets 1, 2, 3) or 5 μM (dataset 4) CytD for 4 h.

To prepare samples for the quantification of luminal filaments, cells were first plated in a 12-well plate at a density of 1 * 10^6^ cells/ well with either 0.1% DMSO or 2 μM TG (final DMSO conc. 0.1%) for 1 h. Cells were then collected, counted and plated on prepared grids (see above) with either 2.5 μM CytD & 0.1% DMSO or 2.5 μM CytD & 2 μM TG. Cells were vitrified 4 h after plating on grids. In total, cells were incubated with CytD for 4 h and with DMSO/ TG for 5 h. The final DMSO concentration in all samples was 0.125%.

For preliminary experiments shown in Fig. EV2A, cells were plated in 2 μM TG or 25 μM MG132 for 12 h. Cells were then collected and plated on grids with 2.5 μM CytD for another 2 h before vitrification. The final DMSO concentration was 0.125% for CytD & TG and 0.275% for CytD & MG132.

Samples from cofilin knock-down cells were prepared on day 7 or 8 of the knock-down. Cells were incubated with 2 μM TG for 3 h, then mixed with 2.5 μM CytD and plated on EM grids following the procedure given above. Samples were vitrified after 5 h of TG and 2 h of CytD treatment.

### Vitrification of specimen

Samples were vitrified similar to the procedure published in Foster *et al*., 2022. Briefly, Whatman No. 1 filter paper stripes were cut to dimensions of approx. 4 × 0.7 cm and the final 0.5 cm were folded over by 90°, generating an L-shaped stripe. Stripes were incubated in a humidified chamber for at least 20 min before use. A Vitrobot MkII (Thermo Fisher) was set to 25°C with 100% humidity and automated blotting was disabled. One grid was taken up with Vitrobot tweezers, placed on the bench and 4 μL of S2 media with 0.1% pluronic-F68 (24040-032, Gibco) and 29% (v/ v) 10 nm BSA-coated gold fiducials in PBS (see below) were carefully placed on the grid. The grid was then loaded onto the Vitrobot, blotted from the backside using the L-shaped filter paper stripes and plunge-frozen in liquid ethane.

BSA-coated gold fiducials were generated by incubating 9.5 mL gold colloid (EM.GC10, BBI solutions) with 0.5 mL 5 mM sodium phosphate buffer, pH 5.0, with 2.5 mg BSA. The final mixture contained 125 μg/ mL BSA and 250 μM sodium phosphate. The sample was incubated at 4°C over-night on a rotating wheel. BSA-coated gold was washed by splitting the volume into 1.5 mL tubes and centrifuging at 25,000 x g for 1 h at 4°C. The supernatant was discarded, pellets were resuspended in PBS and the procedure was repeated. Finally, pelleted BSA-coated gold was pooled and resuspended in a final volume of 200 – 250 μL PBS. Fiducials were stored at 4°C for 6 – 12 months.

### Electron microscopy data acquisition and tomogram reconstruction

Tilt series were acquired on a 300 kV FEI Titan Krios (Thermo Fisher) equipped with a K2 or K3 detector and energy filter (Gatan, slit width 20 eV) using SerialEM software (Mastronarde, 2005). A dose-symmetric scheme (Hagen *et al*., 2017) was used to acquire a tilt series with 2° or 3° increments between ±60°. 10 or 14 dose-fractionated images were acquired at each tilt with a total dose of 112 – 124 e^-^/ Å^2^. Data was acquired with a pixel size of 2.952 or 2.659 Å/ pixel and a nominal defocus of -2.5 to -6 μm. Details for each dataset are in Appendix Tables S3 – 5.

All pre-processing and reconstruction steps were performed in WARP 1.0.9 (Tegunov and Cramer, 2019) unless stated otherwise. Frames were aligned and gain-corrected. The defocus was estimated and bad tilt images were manually removed. A trained model was used to pick fiducials and mask them during tomogram reconstruction. Tilt series were aligned in IMOD (version 4.10.49), AreTomo (version 1.1.1) or in MATLAB using dautoalign (https://github.com/alisterburt/autoalign_dynamo). Tilt series were CTF-corrected and tomograms were reconstructed with a binning of 4 (bin4, 11.81 Å/ pixel or 10.64 Å/ pixel). For quantifications and images shown in figures, bin4 tomograms were deconvolved in WARP using a Wiener-like filter (Tegunov and Cramer, 2019).

Images of slices through tomograms were generated by opening bin4, deconvolved tomograms in IMOD, setting the slice thickness to 5 and adjusting the angles in the slicer window.

### Quantification of protrusion thickness

The thickness of protrusions from vitrified specimen (datasets 1 – 4) was measured in IMOD using the distance measurement tool. Briefly, bin4, deconvolved tomograms (11.81 Å/ pixel) were opened in IMOD and adjusted in the slicer window to show the cross section of the protrusion with 10 overlaid slices. For each protrusion, we estimated the thickness at 3 positions along the protrusion and determined the average protrusion thickness. The graph depicts the average thickness of each protrusion. If two protrusions were present in the same tomogram, they were measured separately. 111 protrusions from 108 tomograms were 150.1 nm ± 38.95 (mean ± SD) thick. Protrusions from different biological replicates are colored differently.

### Subtomogram averaging and classification of microtubules

Microtubules were picked in IMOD from cross-sections of bin4, deconvolved tomograms (11.81 or 10.64 Å/ pixel). Approx. 10 points were placed in the middle of each microtubule along the protrusion. Each microtubule was picked in a different contour. Model points were exported as coordinates and imported into Dynamo 1.1.460 (Castaño-Díez *et al*., 2012) as a filament with torsion model workflow using a custom script, as described in Foster *et al*., 2022. Particles were cropped from bin4 tomograms every 8 nm with a box size of 50 pixels. Particles were averaged and jointly aligned to the generated average. After joint alignment, particles from each microtubule were averaged to generate per-microtubule averages. Across our data (>1,000 microtubules), we found microtubules with 12, 13, 14 and 15 protofilaments. We chose ‘good-looking’ individual filament averages of each protofilament number and rotated each of them by 180° around the axis perpendicular to filament axis to generate a reference with the same protofilament number but opposite polarity. The resulting 8 references (12, 13, 14, 15 protofilaments, each with plus and minus-ends facing towards the plus end of the filament axis) were used for an initial multireference alignment of all particles of one dataset in Dynamo. The class averages after this alignment had higher signal-to-noise ratios than the initial references because more particles were present in each average, and they were used as initial references for all the following multireference alignments.

To automatically determine the polarity and protofilament number of each microtubule, the aligned particle table was first cleaned by retaining 80% of particles with the best cross-correlation scores. A custom MATLAB script was used to assess the percentage of particles in each filament that classified into either class and to assign each filament to a protofilament number and orientation. A microtubule was assigned if at least 65% of particles in that filament classified into that class or if at least 50% of particles classified into a single class and the overall cross-correlation score was equal or higher 0.13. Microtubules that did not match either of these criteria remained undetermined (30 out of 574) and were excluded from all quantifications. The classification and analysis procedures were validated across multiple datasets by comparing the results of this analysis to visual inspection of individually averaged microtubules. Scripts used for the export of IMOD filament models and for the determination of microtubule polarity and protofilament number have been uploaded to GitHub (https://github.com/carterlablmb/Ventura-Rogers-Carter-2023.git).

Using this procedure, we determined the protofilament number of 544 microtubules from 111 protrusions from 3 biological replicates and 5 technical replicates. Technical replicates were pooled and the percentage of 12, 13, 14, 15 protofilament microtubules was determined for each biological replicate as shown in Fig. 1D. Data from all replicates was pooled to show the number of protrusions containing microtubules with different protofilament numbers in Fig. EV1F.

The protofilament numbers of microtubules from DMSO and TG treated cells were determined by classification as described above. We then determined which microtubules contained luminal filaments and which did not. We calculated the percentage of microtubules with 12, 13, 14, 15 protofilaments in each biological replicate for microtubules with or without luminal filaments as shown in Fig. EV2E. For statistical analysis, means and SDs of the normalized protofilament number percentages were calculated for microtubules without (347) and with (82) luminal filaments. Samples treated with DMSO or TG were joined for this analysis as we found no difference in protofilament number distribution (multiple unpaired t-tests, non-significant). Multiple unpaired t-tests were performed to determine if the percentage of each protofilament number was altered and all comparisons showed no significance. The number of microtubules analyzed in each biological replicate under each condition can be found in Appendix Table S3. 35/ 119, 21/ 139 and 30/ 192 microtubules from biological replicates 1 – 3 (datasets 5 – 7), respectively, contained luminal filaments. 21 out of 450 microtubules had unassigned protofilament number and were excluded from the analysis, leading to the following numbers for microtubules with luminal filaments for the three biological replicates: 35/ 117, 18/ 130, 29/ 182.

### Analysis and quantification of luminal filaments

To quantify the occurrence and lengths of luminal filaments, we manually picked them from bin4 deconvolved tomograms in IMOD. Filaments were picked in the slicer window from side views and adjusted in a second slicer window showing a cross section of the same filament. One filament was picked per contour.

To quantify the percentage of microtubules that contained luminal filaments, we mapped each filament to its surrounding microtubule and determined the percentage of microtubules that contained at least one luminal filament (Fig. 2C). An unpaired t-test was used to determined statistical significance between percentages from three biological replicates (technical replicates were pooled). To determine the filament length, we extracted the filament contour length using the ‘imodinfo’ command (Fig. 2D). The filament lengths were not normally distributed as determined from a D’Agostino & Pearson test applied to each biological replicate and each treatment. Thus, we performed a non-parametric Mann-Whitney test and found no statistically significant difference in luminal filament lengths between CytD & DMSO and CytD & TG treated samples (Fig. 2D).

To determine which proportion of microtubule was covered by luminal filaments, we placed multiple points of a single contour along the length of each microtubule and used the ‘imodinfo’ command to extract the contour (i. e. microtubule) length. We then divided the length of the luminal filaments within a single microtubule by the length of the surrounding microtubule (Fig. 2E). A coverage of 1 means that the entire microtubule was covered with a luminal filament. A coverage of 0 means that there were no filaments within this microtubule. If there were multiple filaments within the same microtubule, their lengths were summed prior to division by the microtubule length. 71 out of 206 microtubules from cells treated with CytD & TG contained at least one luminal filament. Of these, the majority (39/ 71, 54.9%) of microtubules showed a coverage of 0.1 – 0.3 with luminal filaments.

To quantify the lengths of luminal filaments with cofilactin or non-cofilactin morphology by visual inspection (Fig. EV3I), we measured the length of each filament and classified it as cofilactin-like or non-cofilactin like. Examples for cofilactin and non-cofilactin morphologies can be found in the right panel of Fig. EV3I. For filaments that transitioned between morphologies, we measured the lengths of each morphology. Percentages of each morphology were determined for three biological replicates (technical replicates were pooled) and mean with SDs are shown in Fig. EV3I. An unpaired t-test was used to determine statistical significance between the proportion of luminal cofilactin filaments in control (90.2 ± 6.8%, mean ± SD) and cofilin dsRNA (25.2 ± 21.8%) knock-down cells.

### Determination of the cross-over distance of filaments from Fourier transformations

Three luminal filaments from well-aligned bin4 tomograms (11.81 Å/ pixel or 10.64 Å/ pixel) were oriented horizontally in the xy-plane using the --rot flag in EMAN2 (Tang *et al*., 2007) or newstack in IMOD. The volumes were then trimmed using IMOD’s trimvol tool to a box size of 200 – 285 pixels and masked with a gaussian cylindrical mask with a diameter of approx. 28 nm to exclude densities outside the luminal filament and the surrounding microtubule. The masked volumes were projected along the filament axis and Fourier transformed in ImageJ. Fourier transforms with a box size of 512 pixels were binned by 2 to a box size of 256 pixels. Layer lines corresponding to the cross-over distance were identified and the distance between the equator and the line was measured using the line tool in Image J. The corresponding frequency was calculated with the following formula: box size [pixel^2^] * pixel size [nm/ pixel] / distance between equator and layer line [pixel].

### Subtomogram averaging of luminal filaments

Microtubules from dTAT knock-out cells contained luminal filaments with similar appearance as those from wild-type cells (Appendix Figure S1C, dataset 8) and this data was joined with datasets 5 – 7 for subtomogram averaging of luminal filaments. All datasets were first processed independently until stated otherwise. IMOD models of luminal filaments picked manually for the quantification of luminal filament lengths (see Analysis and quantification of luminal filaments) were exported into Dynamo ‘filament with torsion’ models using a custom script as described in Foster *et al*., 2022. Particles were cropped at a 4-times binned pixel size (11.81 or 10.64 Å/ pixel) with a box size of 50 pixels and with a particle distance of 6 nm. To generate an initial reference, particles were aligned to 5 references generated by averaging 50 randomly chosen particles in a multireference alignment in Dynamo with a cylindrical soft-edged mask. The cone-search along the filament was limited to 30°, the in-plane angular search to 180° and the shift along the filament axis to 2.4 nm during the alignment with 6 iterations. A good reference was chosen from the final classes and used as initial reference for a joint alignment of all particles.

To obtain a reference for determining the orientation of filaments, particles from each filament were averaged after joint alignment. A good average was selected and rotated by 180° using EMAN2 to obtain a reference with opposite filament polarity. These two references were used in multireference alignments in Dynamo. Filament polarities were determined with a custom-script that assessed the proportion of particles in each filament that classified into each polarity class. Filaments were assigned to the polarity in which the majority of particles classified. Particles from filaments with one polarity were rotated by 180° around the second Euler angle using subTOM software (version 1.1.2) (https://github.com/DustinMorado/subTOM/releases/tag/v1.1.4). This rotation was applied to the original particle positions, not the particle positions obtained after the initial alignment. An array of random subsets was generated by placing particles from each filament into the same subset to avoid duplicate particles in separate groups used for resolution estimation.

To transition to Relion, Dynamo particle tables were converted to Relion starfiles using the dynamo2warp option of the Dynamo2M package (https://github.com/alisterburt/dynamo2m) and additional columns such as filament number and random subset number (see above) were added using the Starparser package (https://github.com/sami-chaaban/starparser). Particles were reconstructed at a pixel size of 5.90 Å/ pixel in WARP with a box size of 120 pixels. All following steps were performed in Relion 3.1 unless stated otherwise. Particles from each dataset were imported with different optics groups and joined using the ‘join star files’ option. The starting particle number was 7,549 particles. To obtain an initial reference, a mask and helical parameters, particles from dataset 8 were refined without helical symmetry using a cylindrical mask. The resulting reference was then refined with a mask generated in Chimera (Pettersen *et al*., 2004) by fitting a cofilactin filament model (PDB 5YU8) into the structure and using the ‘colorzone’ function to generate a map from the averaged luminal filament density without the surrounding microtubule density. The resulting refined average from dataset 8 was used to determine initial helical symmetry parameters. The average was lowpass filtered (60 Å) and, together with the mask generated in Chimera, used in a 3D classification of particles from all datasets (5 – 8) to select good particles. Particles were classified into 6 classes with T = 1 and helical reconstruction and symmetry (initial twist: -162°, initial rise: 29 Å). A selection of 3,801 particles from 4 classes was further refined. The refined particle positions and orientations were used to re-reconstruct particles at a pixel size of 2.95 Å/ pixel with a box size of 256 pixels in WARP. Particles were aligned in a final refinement with T = 6.

To determine the correct pixel size of the final map, we used the generated Drosophila cofilactin model (see Generation of a *Drosophila* cofilactin model). We fit this model into the post-processed and masked map using the option ‘use map simulated from atoms, resolution 17 Å’ in Chimera. The correlation of the fit was noted in a spreadsheet upon adjusting the pixel size of the post-processed map. The correct pixel size of 2.83 Å/ pixel was estimated from plotting the correlation values and estimating the peak by eye. We then ran another post-processing job to determine the FSC and obtain the filtered map.

The final map contains 3,801 particles. The resolution of 16.5 Å was determined with the corrected pixel size. The luminal filament average was lowpass filtered to the determined resolution and sharpened with a B-factor of - 500 Å^2^ in Relion 3.1. The helical symmetry of the luminal filament average (−161.8° twist, 28.5 Å rise) was determined using the relion_helix_toolbox ‘search’ option, with an outer diameter of 120 Å, a rise of 1 – 40 Å and a twist of -100 to -200°.

### Generation of a *Drosophila* cofilactin model

To generate a *Drosophila* cofilactin model, we predicted the structure of *Drosophila* cofilin (Uniprot ID P45594) together with *Drosophila* actin (Act5C, Uniprot ID P10987) using Alphafold 2-Multimer (Evans *et al*., 2022) through a local installation of Colabfold 1.2.0 (Mirdita *et al*., 2022). The actin moiety in the predicted cofilin-actin unit resembled the actin structure from a previous cofilactin filament model made of chicken cofilin and rabbit actin (PDB 5YU8) (Tanaka *et al*., 2018). The prediction of cofilin also matched the chicken cofilin structure from this model, except for a loop that is absent in *Drosophila* cofilin due to its shorter length (148 residues compared to 166 residues in chicken cofilin).

We aligned the predicted cofilin-actin unit with actin from the cofilactin model (PDB 5YU8) using the ‘matchmaker’ tool in Chimera. We assembled a *Drosophila* cofilactin filament by repeating this procedure for 8 consecutive cofilin-actin units and then removing the two cofilin moieties at the pointed end. The resulting model contains 8 actin and 6 cofilin units. We removed the first 6 N-terminal residues (MCDEEV) and residues 42 – 50 (QGVMVGMGQ) of the DNAse loop of actin, as these residues were absent from the previous PDB model 5YU8 on which we based our cofilactin model. We truncated side chains in Phenix version 1.10-2155 (Adams *et al*., 2010) using the phenix.pdbtools option remove=“protein and sidechain”. To validate the *Drosophila* cofilactin model, we first generated a density map from it using the Chimera ‘molmap’ function at a resolution of 17 Å. We then fit this generated map into our subtomogram average structure of the luminal filaments and obtained a score of 0.963 for atoms inside the density.

### Classification of luminal filaments from control and cofilin knock-down samples

To determine the percentages of particles with cofilactin, bare f-actin and other morphologies with subtomogram classification, we manually picked particles and exported them to Dynamo filament with torsion models and subsequently to Relion 3 star files following workflows described in the above sections. The initial particle number was 9,679 from four biological replicates (dataset 9: 4,168 particles, dataset 10: 1,778 particles, dataset 11:2,743 particles, dataset 12: 990 particles), each containing control (6,271 particles across all datasets) and cofilin knock-down (3,408 particles across all datasets) samples.

Particles were jointly 3D classified into 20 classes with helical symmetry (−165° twist, 29 Å rise) to select cofilactin, bare f-actin and other classes. Particles from cofilactin and other classes were pooled while particles from bare f-actin classes were saved separately. We iteratively classified these subsets to obtain classes with different filament polarities and rotated particles from classes with opposite orientation by -180 degrees. To do this, we first converted the star files of opposite-polarity classes into Dynamo tables (dynamo2m package) and then motive lists, rotated particles using the subTOM package, converted the motive lists back into Dynamo tables and then star files (dynamo2m package), and finally replaced the angles columns (AngleRot, AngleTilt, AnglePsi) from the ‘old’ star file with the values determined after rotation using the Starparser package. The uniform orientation of particles was assessed by eye from repeated 3D classification with helical symmetry.

Once particles were uniformly oriented, we initiated classification to determine the percentage of particles in each class. For this, we performed 9 3D classifications with 5, 10 or 20 classes and each with a helical rise of -162°, -165°, -167° and 25 iterations. These values for the helical rise correspond to those determined for cofilactin (−162°) or f-actin (−167°). -165° was included to represent a value in between the two. Particles were aligned to a feature-less, lowpass filtered (60 Å) rod with a cylindrical soft-edged mask. For each classification, we determined by eye which classes resembled cofilactin, bare f-actin or ‘other’ morphologies. An example of classification results, the assigned morphology and the quantification from this individual classification run can be found in Fig. EV3G, H. We used a custom MATLAB script to determine the number of particles from control or cofilin knock-down cells from each biological replicate in each category of classes (cofilactin-, bare f-actin, other). From this, we calculated the percentages for each biological replicate and control/ cofilin knock-down samples for each classification run. The percentages were averaged (SDs were in between 0.7 – 14.4%) for each biological replicate and are shown as circles (control dsRNA) or triangles (cofilin dsRNA) in Fig. 3L. Unpaired t-tests were used to compare results from classification into cofilactin, bare f-actin and other classes between control and cofilin knock-down samples. Individual class averages representing cofilactin, bare f-actin and other morphologies are shown in the top panel of Fig. 3L.

## Acknowledgements

We than the MRC LMB Electron Microscopy Facility, Scientific Computing and Light Microscopy Facility for access and support. We thank Tom Dendooven and Alister Burt for support with subtomogram averaging, Emmanuel Derivery and Alice Bittlestone for assistance with S2 cell culture. We acknowledge Anna Marie Sokac and Buzz Baum for the gifts of the *Drosophila* cofilin and P-cofilin antibodies. We thank Simon Bullock and members of the Carter laboratory for discussions.

This study was supported by the Medical Research Council (MRC_UP_A025_1011) and Wellcome Trust (210711/Z/18/Z) grants to A.P.C., by the National Institute of Neurological Disorders and Stroke grant (R21NS125795) to S.L.R. and by a PhD studentship from the Cambridge Trust to C.V.S.

## Author contributions

**Camilla Ventura Santos**: Conceptualization, Data curation, Formal analysis, Investigation, Methodology, Validation, Visualization, Writing. **Stephen L. Rogers**: Conceptualization, Resources, Writing. **Andrew P. Carter**: Conceptualization, Funding acquisition, Supervision, Writing.

## Data availability

All raw data and filtered, bin4 tomograms have been uploaded to EMPIAR with the following accession codes: EMPIAR-11450 for Datasets 1 – 4, EMPIAR-11451 for Datasets 5 – 7, EMPIAR-11452 for Dataset 8, EMPIAR-11453 for Datasets 9 – 12.

Representative tomograms have been deposited to the EMDB. EMD-16685, EMD-16693 are examples of tomograms used for the analysis of microtubules in Drosophila protrusions. EMD-16720 shows an exemplary tomogram of an S2 cell protrusion with luminal filaments after treatment with CytD & TG. EMD-16695 is an example tomogram from Drosophila dTAT KO cells used for the subtomogram averaging of luminal filaments. EMD-16800 and EMD-16811 are example tomograms from cofilin knock-down cells.

The subtomogram average structure of the luminal filament and a model of *Drosophila* cofilactin have been deposited to the EMDB (EMD-16877) and PDB (8OH4).

Scripts used in all analyses have been uploaded to GitHub (https://github.com/carterlablmb/Ventura-Rogers-Carter-2023.git). Analysis spreadsheets, western blots and additional files have been deposited at Biostudies (S-BSST1048) (Sarkans *et al*., 2018).

## Expanded View figures

**Figure EV1.**
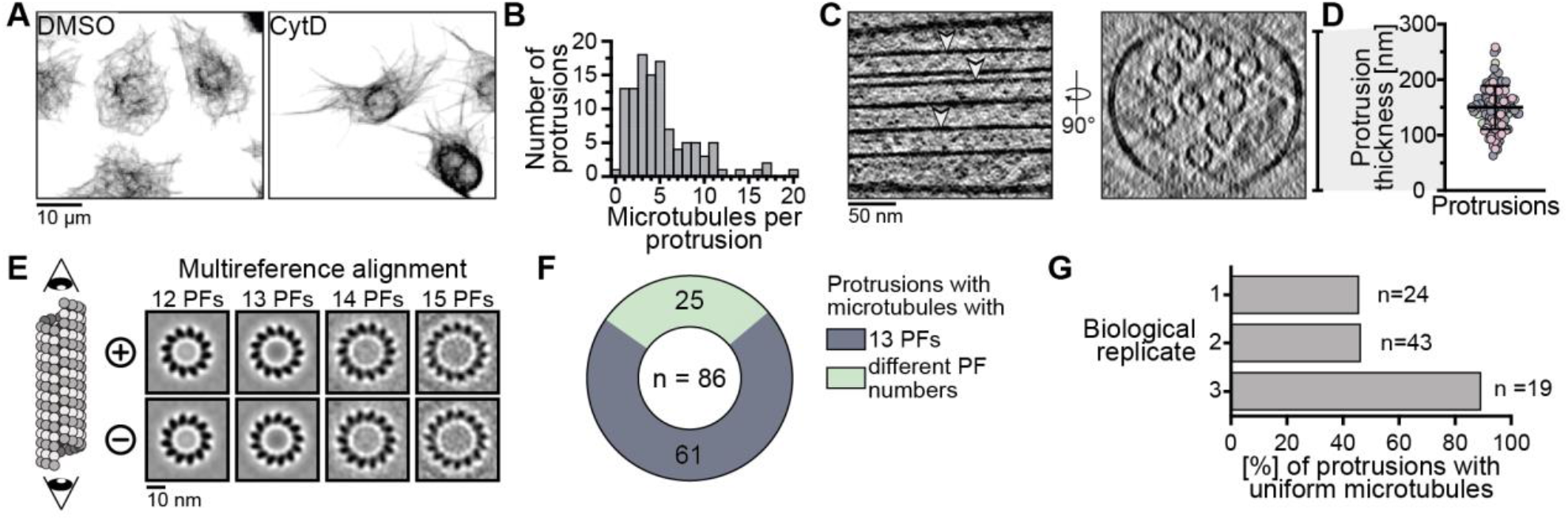
Analysis of microtubules in induced S2 cell protrusions. A) Immunofluorescence staining of α-tubulin in S2 cells shows that CytD treatment induces the formation of protrusions which are filled with microtubules. Control cells treated with a vehicle (DMSO) are shown on the left. B) Histogram depicting the number of microtubules found in protrusions that were targeted for the acquisition of tilt series. C) Tomogram slices of a protrusion (left) and its corresponding cross section (right) showing that microtubules form a parallel array. Arrowheads point at microtubules. The bar on the right shows the measured thickness of the protrusion at this tomographic slice and is exemplary for thickness measurements shown in D). D) Protrusion thicknesses (150.1 ± 39.0 nm, mean ± SD) measured from cross sections of tomograms. Each protrusion was measured at three positions in the tomogram and the average thickness per protrusion is shown. Protrusions from different biological replicates are colored grey, pink and green. E) Cartoon (left) and projections of references (right) used for multireference alignment to determine microtubule protofilament (PF) number and orientation. Protofilaments of microtubules viewed from the plus end appear to rotate in an anti-clockwise direction (top row) (Sosa and Chrétien, 1998). Conversely, they appear to rotate in a clockwise direction when viewed from the minus end (bottom row). Polarities and protofilament numbers were determined based on the proportion of particles inside each microtubule that classified into each of the 8 classes in one iteration of a multireference alignment (for details see Materials and methods). F) Pie chart showing that the majority but not all protrusions contained exclusively 13 protofilament microtubules. We found 25 protrusions that contained microtubules with two or three different protofilament numbers. G) Percentages of protrusions containing uniformly oriented microtubules from three biological replicates show that microtubule uniformity can be variable across datasets.

**Figure EV2.**
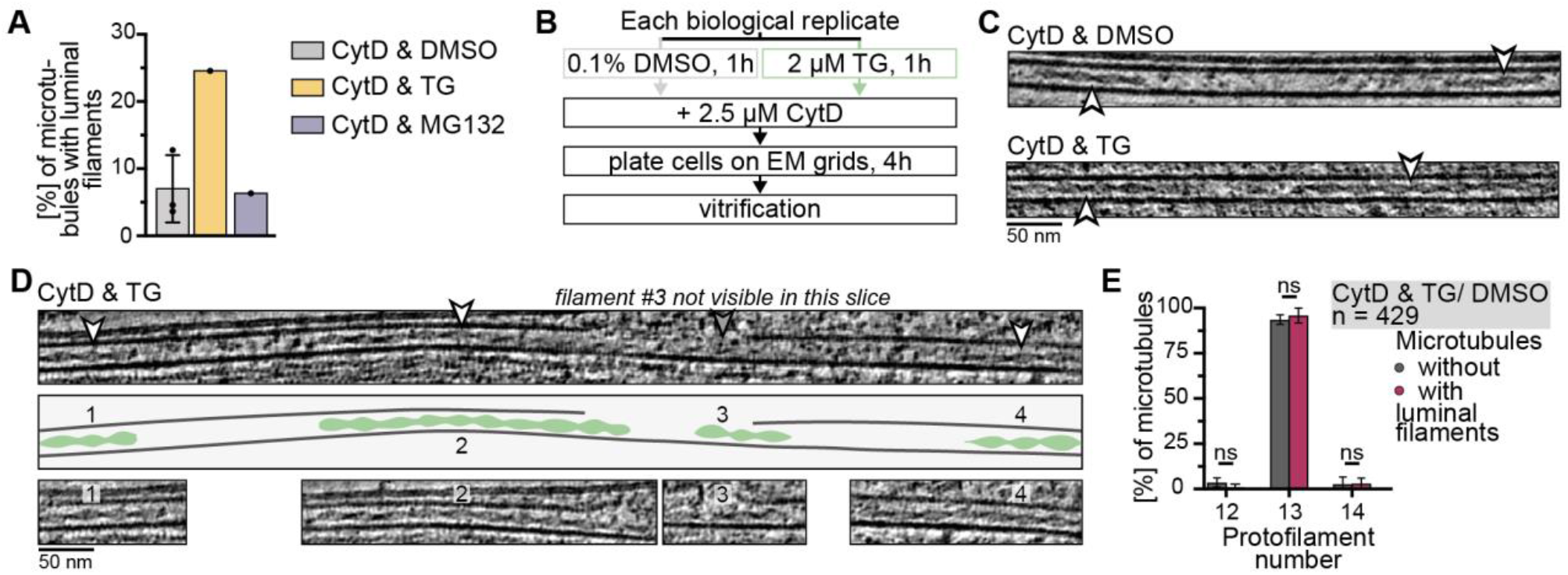
Analysis of filaments inside the microtubule lumen. A) Percentage of microtubules with at least one luminal filament after treatment with CytD & TG (24.6%, mean) or CytD & MG132 (6.4%) in a preliminary experiment. Cells were treated with TG or MG132 over-night before addition of CytD for 2 h followed by vitrification. The control condition (4 h CytD & 5 h DMSO treatment, CytD & DMSO, 7.0 ± 5.0%, mean ± SD) from Fig. 2C is shown for comparison. B) Workflow for the preparation of biological replicates that were treated with CytD & DMSO/ TG and used for the analysis of luminal filaments. C) Tomogram slices showing examples of microtubules with two luminal filaments (white arrowheads) from samples with indicated treatments. D) Tomogram slices and cartoon of a microtubule that contains four luminal filament segments. Top panel shows an overview image of the microtubule where three out of four luminal filaments are visible (white arrowheads). The fourth filament is not clear in this tomographic slice (grey arrowhead). Middle panel shows a cartoon of the microtubule and luminal filaments. The four luminal filaments are also depicted on tomographic slices in the bottom panel with numbers corresponding to those in the cartoon. The example is from cells treated with CytD & TG. E) Percentage of microtubules without (grey) or with (red) luminal filaments that have 12 (3.6 ± 2.6% without, 1.0 ± 1.7% with, mean ± SD), 13 (93.6 ± 2.6% without, 96.0 ± 4.3 with) or 14 (2.8 ± 3.8% without, 3.1 ± 2.9% with) protofilaments. Microtubules from CytD & DMSO and from CytD & TG treated cells were pooled for this analysis. Percentages from three biological replicates are shown and were used for a 2-way ANOVA test with comparisons between microtubules of each protofilament number with and without luminal filaments (12: p = 0.63, 13: p = 0.92, 14: p > 0.99, all ns).

**Figure EV3.**
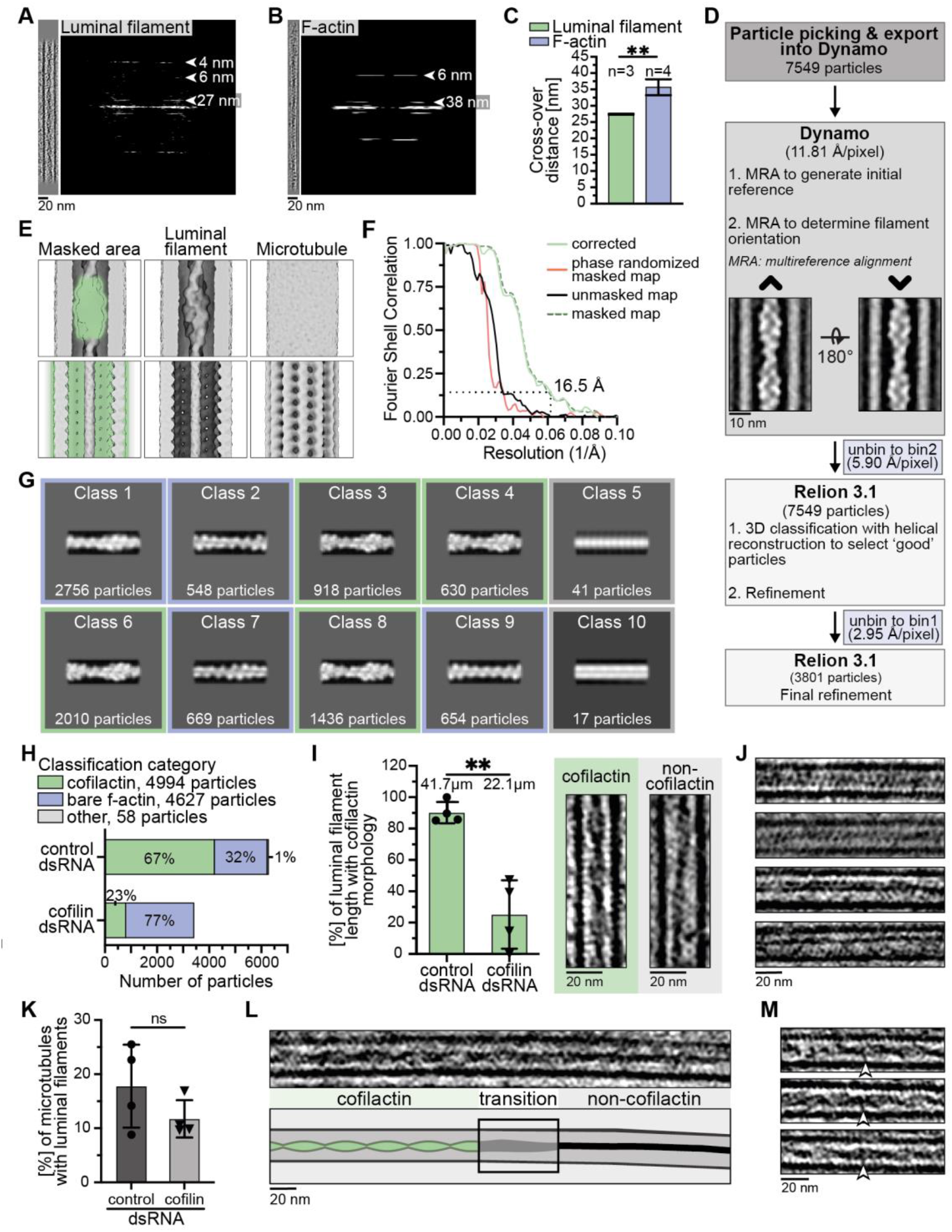
Structural analysis of luminal filaments. A, B) Fourier transforms of the luminal and f-actin filament shown in Fig. 3A, B. Projections of aligned and masked volumes of the filaments used to generate the Fourier transform are shown in the left panels. Layer lines and their corresponding frequencies are indicated with white arrowheads. C) Measurements of cross-over distances of luminal filaments (27.4 ± 0.14 nm, mean ± SD) and cytoplasmic f-actin (35.7 ± 2.4 nm) from Fourier transforms. Statistical significance was assessed with an unpaired t-test (p = 0.0022, **). D) Workflow used for subtomogram averaging of luminal filaments. E) Images showing that the luminal filaments and the surrounding microtubule are not regularly positioned with respect to each other. The top row shows the result of a subtomogram alignment in which the luminal density was included in the mask (green area, left). The luminal filament is well defined (middle) whereas the microtubule density has smeared out (right). The bottom row shows the result of an alignment in which the masked area included the microtubule density but not the luminal filament (green area, left). This alignment resulted in a featureless rod representing the luminal density (middle) and a well-defined microtubule density (right). F) Fourier shell correlation (FSC) of the subtomogram average structure shown in Fig. 3C, D. The corrected (green, continuous) curve crosses the 0.143 criterium at ∼0.061 1/ Å, corresponding to a resolution of 16.5 Å. G) Representative results from a subtomogram classification of luminal filaments from control and cofilin knock-down cells. Legend as in H. Classes assigned to cofilactin, bare f-actin and other morphologies are marked with green, blue and grey borders, respectively. H) Exemplary quantification of the classification shown in H. This quantification was performed for all classifications and averages of the percentages of particles in each assigned morphology (cofilactin, bare f-actin or other) for each biological replicate are shown in Fig. 3L. I) Percentage of luminal filament length resembling cofilactin morphology (90.2 ± 6.8% control dsRNA, 25.22 ± 21.8% cofilin dsRNA, mean ± SD) assessed from visual inspection of the tomograms. The panel on the right shows examples of cofilactin (green) and non-cofilactin (grey) morphologies. We sampled 41.7 μm luminal filament length in control and 22.1 μm in cofilin knock-down cells. Cofilin knock-down cells had significantly less cofilactin-like luminal filaments (unpaired t-test, p=0.0013, **). J) Tomogram slices of luminal filaments from cofilin knock-down cells showing examples of filament morphologies that do not resemble cofilactin or bare f-actin. K) Percentage of microtubules containing at least one luminal filament in samples from control (17.8 ± 7.7%, mean ± SD) or cofilin (11.7 ± 3.4%) knock-down cells. The reduction in cofilin knock-down samples was non-significant (unpaired t-test, p = 0.2011, ns). L) Tomogram slice and cartoon showing a luminal filament that transitions from a cofilactin (left) to a non-cofilactin (right) morphology. M) Examples of linking densities (white arrowheads) between luminal filaments and the surrounding microtubule wall.

### Appendix

**Appendix Table S1.**
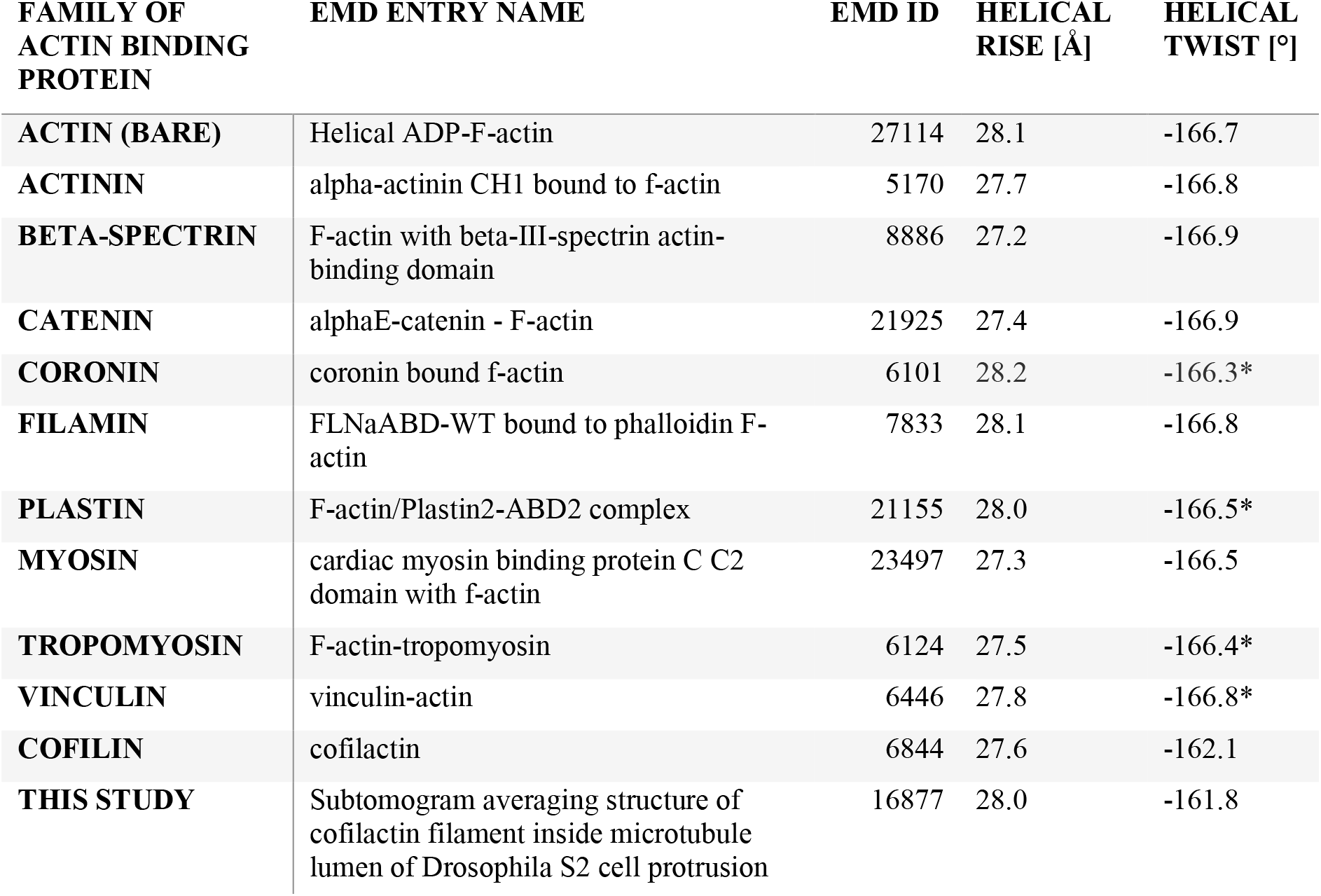
EMDB entries of actin together with actin-binding proteins solved by helical reconstruction. Helical twist values marked with a star were deposited as positive values but are displayed with a negative sign because they describe left-handed helix symmetries. Helical parameters were rounded to one decimal.

| **FAMILY OF ACTIN BINDING PROTEIN** | **EMD ENTRY NAME** | **EMD ID** | **HELICAL RISE [Å]** | **HELICAL TWIST [°]** |
| --- | --- | --- | --- | --- |
| **ACTIN (BARE)** | Helical ADP-F-actin | 27114 | 28.1 | -166.7 |
| **ACTININ** | alpha-actinin CH1 bound to f-actin | 5170 | 27.7 | -166.8 |
| **BETA-SPECTRIN** | F-actin with beta-III-spectrin actin-binding domain | 8886 | 27.2 | -166.9 |
| **CATENIN** | alphaE-catenin - F-actin | 21925 | 27.4 | -166.9 |
| **CORONIN** | coronin bound f-actin | 6101 | 28.2 | -166.3* |
| **FILAMIN** | FLNaABD-WT bound to phalloidin F-actin | 7833 | 28.1 | -166.8 |
| **PLASTIN** | F-actin/Plastin2-ABD2 complex | 21155 | 28.0 | -166.5* |
| **MYOSIN** | cardiac myosin binding protein C C2 domain with f-actin | 23497 | 27.3 | -166.5 |
| **TROPOMYOSIN** | F-actin-tropomyosin | 6124 | 27.5 | -166.4* |
| **VINCULIN** | vinculin-actin | 6446 | 27.8 | -166.8* |
| **COFILIN** | cofilactin | 6844 | 27.6 | -162.1 |
| **THIS STUDY** | Subtomogram averaging structure of cofilactin filament inside microtubule lumen of Drosophila S2 cell protrusion | 16877 | 28.0 | -161.8 |

**Appendix Figure S2.**
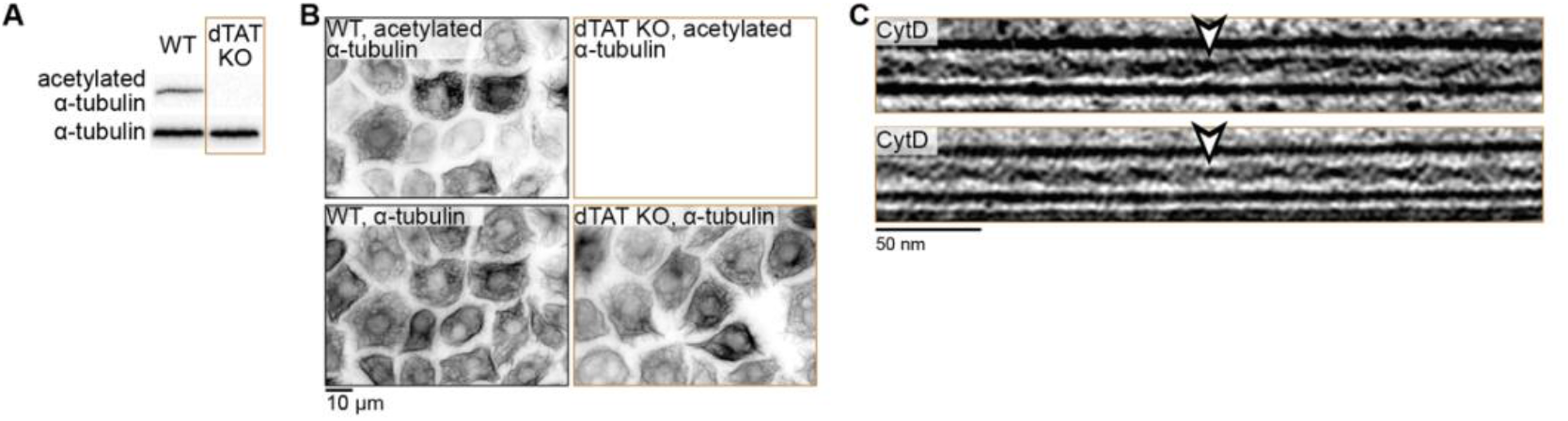
Validation of the *Drosophila* α-tubulin acetylase (dTAT) CRISPR knock-out. A) Western blot of wild-type (WT) or dTAT knock-out (KO) cells blotted for acetylated α-tubulin and α-tubulin (loading control). B) Immunofluorescence of WT (left) and dTAT KO (right) cells stained for acetylated α-tubulin (top) or α-tubulin (bottom). Tomogram slices of luminal filaments (white arrowheads) in dTAT KO cells showing that they have similar appearance to luminal filaments found in WT cells (Fig. 2A, B). dTAT KO cells were treated with 5 μM CytD for 4 h before vitrification.

**Appendix Table S3.** Acquisition and processing parameters for datasets 1 – 4 (EMPIAR-11450). These datasets were used for quantification and subtomogram averaging of microtubules in S2 cells (Fig. 1).

| **MICROTUBULE**<br>**ANALYSIS** | **BIOLOGICAL**<br>**REPLICATE 1** | **BIOLOGICAL REPLICATE**<br>**2** |  | **BIOLOGICAL**<br>**REPLICATE 3** |
| --- | --- | --- | --- | --- |
| **GRID ID** | **DZ4** | **DY2** | **DZ1** | **FB3/ FB4** |
| **DATASET** | **dataset 1** | **dataset 2** | **dataset 3** | **dataset 4** |
| **GENOTYPE** | WT | WT |  | WT |
| **SAMPLE CONDITIONS** | 4 h 2.5 μM CytD | 4 h 2.5 μM CytD | | 4 h 5 μM CytD |
| **INCREMENTS/**<br>**NUMBER OF TILTS** | 2°/ 61 | 2°/ 61 | 3°/ 41 | 3°/ 41 |
| **FRAMES PER TILT**<br>**IMAGE** | 10 | 10 | 14 | 14 |
| **PIXEL SIZE [$\text{\AA}$/ PIXEL],**<br>**DETECTOR** | 2.952, K2 | 2.952, K2 |  | 2.952, K2 |
| **TOTAL DOSE [$e^-/\text{\AA}^2$]** | 123.44 | 121.68 | 123.57 | 122.25 |
| **TILT ALIGNMENT**<br>**PROGRAM** | dautoalign and IMOD | IMOD |  | IMOD |
| **ACQUIRED TILT**<br>**SERIES** | 35 | 22 | 39 | 24 |
| **ANALYSED**<br>**TOMOGRAMS** | 33 | 18 | 36 | 22 |
| **NUMBER OF**<br>**PROTRUSIONS** | 33 | 56 |  | 22 |
| **NUMBER OF**<br>**MICROTUBULES** | 109 | 299 |  | 166 |

**Appendix Table S4.** Acquisition and processing parameters for datasets 5 - 7 (EMPIAR-11451) and dataset (EMPIAR-11452). Datasets 5 – 7 were used for quantification of luminal filaments (Fig. 2). Datasets 5 – 8 were used for subtomogram averaging of luminal filaments (Fig. 3). Dataset 8 was collected on the dTAT knock-out mutant.

| *CYTD & DMSO AND CYTD & TG* | BIOLOGICAL REPLICATE 1 |  | BIOLOGICAL REPLICATE 2 |  | BIOLOGICAL REPLICATE 3 |  | DTAT KNOCK-OUT |
| --- | --- | --- | --- | --- | --- | --- | --- |
| ACQUISITION DATE/<br>GRID ID | 17.06.2022 |  | 20.07.2022 |  | 21.07.2021 |  | FE2/ FE4 |
| DATASET | dataset 5 |  | dataset 6 |  | dataset 7 |  | dataset 8 |
| GENOTYPE | WT |  | WT |  | WT |  | dTAT KO |
| SAMPLE CONDITIONS | 5 h 0.1% DMSO,<br>4 h 2.5 µM CytD,<br>0.125% total DMSO | 5 h 2 µM TG,<br>4 h 2.5 µM CytD,<br>0.125% total DMSO | 5 h 0.1% DMSO,<br>4 h 2.5 µM CytD,<br>0.125% total DMSO | 5 h 2 µM TG,<br>4 h 2.5 µM CytD,<br>0.125% total DMSO | 5 h 0.1% DMSO,<br>4 h 2.5 µM CytD,<br>0.125% total DMSO | 5 h 2 µM TG,<br>4 h 2.5 µM CytD,<br>0.125% total DMSO | 4 h 5 µM CytD |
| INCREMENT S/ NUMBER OF TILTS | 3°/ 41 |  | 3°/ 41 |  | 3°/ 41 |  | 3°/ 41 |
| FRAMES PER TILT IMAGE | 14 |  | 14 |  | 14 |  | 14 |
| PIXEL SIZE [Å/ PIXEL], DETECTOR | 2.952, K2 |  | 2.952, K2 |  | 2.659, K3 |  | 2.952, K2 |
| TOTAL DOSE [e/ Å<sup>2</sup>] | 118.08 |  | 122.19 |  | 121.54 |  | 122.25 |
| TILT ALIGNMENT PROGRAM | IMOD |  | AreTomo |  | AreTomo |  | IMOD |
| ACQUIRED TILT SERIES | 12 | 14 | 23 | 18 | 25 | 18 | 65 |
| ANALYSED TOMOGRAMS | 12 | 12 | 19 | 14 | 20 | 13 | 62 |
| NUMBER OF ANALYSED PROTRUSIONS | 10 | 14 | 18 | 15 | 17 | 13 | 62 |
| NUMBER OF MICROTUBULES | 119 |  | 139 |  | 192 |  | 447 |

**Appendix Table S5.** Acquisition and processing parameters for datasets 9 – 12 (EMPIAR-11453). These datasets were used for quantification of luminal filaments upon control or cofilin knock-down and treatment with CytD & TG (Fig. 3J – L). 8 tilt series from biological replicate 1 (dataset 11) and 3 tilt series from replicate 3 (dataset 13) were collected in different acquisition settings.

| **CONTROL AND COFILIN KNOCK-DOWN** | **BIOLOGICAL REPLICATE 1** |  | **BIOLOGICAL REPLICATE 2** |  | **BIOLOGICAL REPLICATE 3** |  | **BIOLOGICAL REPLICATE 4** |  |
| --- | --- | --- | --- | --- | --- | --- | --- | --- |
| **ACQUISITION DATE/ GRID ID** | **08.12.2022 (part 1) & 17.12.2022 (part 2)** |  | **16.12.2022** |  | **02.02.2023 (part 1) & 12.12.2022 (part 2)** |  | **03.02.2023** |  |
| **DATASET** | **dataset 9** |  | **dataset 10** |  | **dataset 11** |  | **dataset 12** |  |
| **DAY 1 OF KNOCK-DOWN** | 01.12.2022 |  | 09.12.2022 |  | 24.11.2022 |  | 17.01.2023 |  |
| **GENOTYPE** | Control knock-down | Cofilin knock-down | Control knock-down | Cofilin knock-down | Control knock-down | Cofilin knock-down | Control knock-down | Cofilin knock-down |
| **SAMPLE CONDITIONS** | 5 h 2 μM TG, 2 h 2.5 μM CytD. 0.125% total DMSO | | 5 h 2 μM TG, 2 h 2.5 μM CytD. 0.125% total DMSO | | 5 h 2 μM TG, 2 h 2.5 μM CytD. 0.125% total DMSO | | 5 h 2 μM TG, 2 h 2.5 μM CytD. 0.125% total DMSO | |
| **INCREMENTS/ NUMBER OF TILTS** | 3°/ 41 |  | 3°/ 41 |  | 3°/ 41 |  | 3°/ 41 |  |
| **FRAMES PER TILT IMAGE** | 14 |  | 14 |  | 14 |  | 14 |  |
| **PIXEL SIZE [$\text{\AA}$/ PIXEL], DETECTOR** | 2.659, K3 (08.12.2022) & 2.952, K2 (17.12.2022) |  | 2.952, K2 |  | 2.952, K2 (02.02.2023) & 2.659, K3 (12.12.2022) |  | 2.952, K2 |  |
| **TOTAL DOSE [$\text{e}/ \text{\AA}^2$]** | 113.74 (08.12.2022) & 111.61 (17.12.2022) |  | 111.61 |  | 119.66 (part 1) & 111.61 (part 2) |  | 120.91 |  |
| **TILT ALIGNMENT PROGRAM** | IMOD & AreTomo |  | AreTomo |  | IMOD |  | IMOD |  |
| **ACQUIRED TILT SERIES** | 21 | 20 | 19 | 19 | 22 | 15 | 14 | 13 |
| **ANALYSED TOMOGRAMS** | 19 | 17 | 16 | 18 | 22 | 13 | 14 | 13 |
| **NUMBER OF ANALYSED PROTRUSIONS** | 20 | 17 | 18 | 18 | 22 | 14 | 14 | 13 |

